# Delayed differentiation of vaginal and uterine microbiomes in dairy cows developing postpartum endometritis

**DOI:** 10.1101/365346

**Authors:** Raúl Miranda-CasoLuengo, Junnan Lu, Erin J. Williams, Aleksandra A. Miranda-CasoLuengo, Stephen D. Carrington, Alexander C.O. Evans, Wim G. Meijer

## Abstract

Bacterial infection of the uterus is a normal event after parturition. While the healthy cow achieves uterine clearance early postpartum, cows unable to control the infection within 21 days after calving develop postpartum endometritis. Studies on the Microbial Ecology of the bovine reproductive tract have focused on either vaginal or uterine microbiomes. This is the first study that compares both microbiomes in the same animals. Terminal Restriction Fragment Length Polymorphism of the 16S rRNA gene showed that despite large differences associated to individuals, a shared community exist in vagina and uterus during the postpartum period. The largest changes associated with development of endometritis were observed at 7 days postpartum, a time when vaginal and uterine microbiomes were most similar. 16S rRNA Pyrosequencing of the vaginal microbiome at 7 days postpartum showed at least three different microbiome types that were associated with postpartum endometritis. All three microbiome types featured reduced bacterial diversity. Taken together, the above findings support a scenario where disruption of the compartmentalization of the reproductive tract during parturition results in the dispersal and mixing of the vaginal and uterine microbiomes, which subsequently are subject to differentiation. This microbial succession is likely associated to early clearance in the healthy cow. In contrast, loss of bacterial diversity and dominance of the microbiome by few bacterial taxa were related to a delayed succession in cows developing endometritis at 7 DPP.

## Introduction

Uterine infection is a common event in the postpartum period in cattle [1]. Early postpartum endometrial inflammation has been shown to occur in response to infection and tissue damage, and as a pre-requisite for uterine clearance and involution in preparation for a future pregnancy [2,3]. Failure or delay in resolving infection has implications for the reproductive health and fertility of cows. Postpartum endometritis is clinically defined as a persistent uterine infection with purulent discharge beyond 21 days postpartum (DPP) and is a major cause of infertility and economic loss in the dairy industry worldwide [4]. A scoring system based on the characterization of vaginal mucus, which reflects the level of bacterial infection in the uterus, has been used as a diagnostic tool for postpartum endometritis [5].

Early studies to identify possible aetiological agent(s) of postpartum uterine infection were focused on the isolation of bacteria from diseased animals [5–8]. Although postpartum endometritis often results from non-specific infections [8], the most common pathogens associated with the uterus of endometritic animals are *Escherichia coli*, *Trueperella pyogenes*, *Prevotella melaninogenicus* and *Fusobacterium necrophorum* [5,9]. These bacteria are often associated in mixed infections of the uterus, with evidence pointing to a succession in which *E. coli* is most prevalent in metritic cows during the first week postpartum. Its presence then increases the subsequent risk of infection by *T. pyogenes* in postpartum weeks 2 and 3, which in turn has been associated with postpartum endometritis [9,10].

A deeper insight into the bovine uterine microbiome at the postpartum period was recently gained using cultivation-independent molecular techniques [11–18]. Temporal analysis showed the occurrence of a bacterial succession in the uterine microbiome, with changes in the composition of cows with uterine disease reported from calving until late postpartum [12,15,17]. The vaginal microbiome has become the subject of analysis using culture-dependent and culture-independent approaches [19–24]. However, a missing piece in the microbiology of the postpartum period is the comparison of the vaginal and uterine microbiomes. This study addresses whether there are consistently distinct communities in these compartments of the reproductive tract and how they are affected by postpartum disease, specifically by postpartum endometritis. In addition, given the differential ability to clear postpartum uterine infection, we hypothesize that the disruption of the compartmentalization in the reproductive tract during parturition results in the mixing of vaginal and uterine microbiota and that differentiation of microbial communities between vagina and uterus will differ between healthy cows and cows developing postpartum endometritis.

## Materials and Methods

### Animals

Three Irish dairy farms participated in this study. Each farm was selected based on high animal welfare standards. From the three farms, 113 cows were recruited onto the study: 42 from Farm A, 37 from Farm B and 34 from Farm C. Sixteen cows were excluded from the study for the following reasons: caesarean operation (1), twin birth (4) uterine wash immediately after calving (1), died (5), sold (3) and dried off (2). A total of 97 cows were included in the study. All procedures were carried out under authorisation of the Irish Department of Health and Children in compliance with the Cruelty to Animals Act 1876 (as amended by EU directive 86/609/EC), and all experimental protocols were approved by the University College Dublin Animal Research Ethics Committee (AREC-P-10-53).

### Vaginal mucus assessment and uterine health diagnosis

Uterine health was assessed by weekly vaginal mucus scoring [5]. The vulva was thoroughly cleaned by spraying a solution of Hibiscrub (BCM Ltd, UK) and subsequent drying with a paper towel (Wypall L35; Kimberley Clark, UK). A clean, lubricated, gloved-hand was then inserted through the vulva. In each cow, the lateral, dorsal and ventral walls of the vagina and the external cervical os were palpated, and the mucus contents of the vagina withdrawn manually for examination. The vaginal mucus was assessed for colour, proportion and volume of pus, and a character score assigned as follows: 0 = clear or translucent mucus; 1 = mucus containing flecks of white or off-white pus; 2 = < 50 ml exudate containing ≤ 50% white or off-white mucopurulent material; and 3 = > 50 ml exudate containing purulent material, usually white or yellow, but occasionally sanguineous. The vaginal mucus was also assessed by odour, and given a score 0 for normal odour or a score of 1 if a fetid odour was detected. Animals were assessed weekly from 7 to 21 DPP and again prior to first breeding (at 50 DPP). They were diagnosed as healthy if the vaginal mucus character score was 0 or 1 and there was no fetid odour present at every time point. Animals were diagnosed as having postpartum endometritis if the vaginal mucus character score was 0 or 1 and there was no fetid odour present on Days 7 and 14 postpartum but then subsequently had purulent mucus i.e. character score of 2 or 3 with or without a fetid odour, on day 21 postpartum or prior to breeding [25].

### Uterine and vaginal swab collection

A double-guarded instrument (Labstock, Co. Meath, Ireland) was used to collect uterine and vaginal swabs in duplicate from each cow. Uterine swabs were collected from the uterine body using a validated method [5,26]. Briefly, the vulva was thoroughly cleaned as described above. A double guarded instrument containing the swab was then inserted through the vagina and cervical canal into the lumen of the uterus, guided by palpation per rectum. Within the uterine body, the cotton swab was extruded from the double guard tube and brought into firm contact with the endometrium by gentle pressure per rectum, about 2cm from the bifurcation of the horns, before being withdrawn into the guard. Vaginal swabs were collected using a similar protocol. Briefly, the double guarded instrument was inserted into the vagina, the swab was extruded from the guard and rotated gently against the vaginal wall before being drawn back into the guard. Swab samples were collected before vaginal examination and mucus assessment to avoid the possibility of introducing bacterial contaminants into the vagina and/or disrupting the equilibrium of the reproductive tract prior to sample collection.

The tip of each swab was cut off and placed into a 1.5 ml polypropylene tube containing 300 µl of TE buffer (20 mM TrisHCl, pH 8.0, 2 mM sodium EDTA), snap-frozen in liquid Nitrogen and shipped in dry ice for molecular analysis.

### DNA extraction

Swab samples were vortexed to disperse cells from the cotton tip and centrifugated for 8 min at 8000 × *g*. Metagenomic DNA was extracted using the Qiagen DNeasy Blood and Tissue kit following the manufacturer’s instructions for Gram positive bacteria (Qiagen). Briefly, pellets were suspended by vortexing in 90 µl TET buffer (TE supplemented with 0.2% [v/v] TritonX-100). 90 µl of TET supplemented with 40 mg/ml egg white Lysozyme were added and incubated at 37°C for 2h. Proteinase K digestion of the sample was performed at 56°C for 1h. Samples were further incubated at 90°C for 5 min and after adding AL buffer, they were loaded in the Qiagen column. Elution was performed in 50 µl of AE buffer. Extracted DNA was stored at −80°C until further use.

### Terminal-Restriction Fragment Length Polymorphism

Terminal-restriction fragment length polymorphism (T-RFLP) was used to obtain fingerprints of the microbial communities associated with vaginal and uterine DNA samples as previously described [27]. Briefly, amplicons of the 16S rRNA genes were obtained by nested PCR, first by 15 amplification cycle with primers 27F-CM (5’-AGAGTTTGATCMTGGCTCAG) and 1492R (5’-TACGGYTACCTTGTTACGACTT) [28,29], followed by a second amplification of a ~1kb fluorescently-labelled product using primers 6FAM-27F-CM and U1052R (5’-GARCTGRCGRCRRCCATGCA) [30]. *Msp*I digested products were ethanol precipitated, resuspended in Hi-Di Formamide (final concentration 50 ng/μl) containing GeneScan-500 LIZ Size Standard (Applied Biosystems) and separated by capillary electrophoresis using a 3130*xl* capillary array (36 cm) in an ABI 3130*xl* Genetic Analyzer (Applied Biosystems).

### Pyrosequencing

A fragment of ~507 bp covering V1-V3 regions of the 16S rRNA gene (*E. coli* position 27 to 534) was selected as target for pyrosequencing. Libraries for pyrosequencing were obtained by nested PCR. In a first step, ~1.5 kb amplicons (10 µL) were produced from each DNA sample with primers 27F-CM and 1492R using the Phusion HF DNA polymerase (New England Biolabs). Then, 1 µL of the resulting amplicon was used as template for the nested PCR (20 µL) using primers A-MID-27F (5’<u>CCATCTCATCCCTGCGTGTCTCCGACTCAG</u>NNNNNNNNNNAGAGTTTGATCMTGGCTCAG) and B-534R (5’<u>CCTATCCCCTGTGTGCCTTGGCAGTCTCAG</u>ATTACCGCGGCTGCTGG). Amplicons were barcoded by introducing a 10 bp multiplex identifier (MID) represented by Ns in the primer A-MID-27F. Underlined sequences at the 5’ of the primers correspond to the pyrosequencing adapters A and B, respectively (454 Life Sciences). PCR reactions contained 1X Phusion HF buffer, 0.2 mM dNTPs, 0.5 µM each of primer (27F-CM and 1492R), 3% DMSO and 0.2 units of Phusion HF DNA polymerase as recommended by the manufacturer (New England Biolabs). The amplification program consisted of an initial denaturation step at 98°C 30 sec, followed by either 15 or 27 cycles of 8 sec melting at 98°C; 20 sec of annealing at 65°C and 45 sec of extension at 72°C for the first and second PCR, respectively. A final extension was carried out for 5 minutes at 72°C. Duplicate PCR amplicons originated from the same sample were pooled and purified with the Agencourt AMPure XP PCR Purification system following the manufacturer’s instructions (Agencourt Bioscience Corporation, Beckman Coulter). The quantification of purified PCR amplicons was assessed in black 96-well plates on a Varioskan spectrofluorometer (Thermo Electron Corporation) using PicoGreen dsDNA Quantitation Kit (Invitrogen). Purified amplicons (1 µL) were also visualized on 2% agarose gel. Equimolar amounts of amplicons obtained from different samples were pooled. Emulsion PCR and 454 library generation were performed at the 454 Sequencing Centre (Branford, USA). Sequencing, starting from the A adapter end by using lib-L annealing beads, was performed on a Roche/454 GS-FLX Titanium system at the 454 Sequencing Centre (Branford, USA).

### Bioinformatics

T-RFLP fragment sizes were determined using GeneMapper v4.0 (Applied Biosystems). Merging of biological replicates and multiple alignment of T-RFLP profiles was performed with T-Align [31]. Only fragments present in both biological replicates and contributing at least 0.5% of the total fluorescence signal were included in the analysis. Terminal restriction fragments and Bray-Curtis resemblance matrix are provided as supporting information (Table S1). Heatmaps of relative abundance obtained from the fluorescent signal of terminal restriction fragments were made using the conditional formatting tool in Excel 2010. Cells were formatted depending on their value using a 3-colour scale where the midpoint was set as the 95 percentile. Data were analysed using Primer6 v6.1.13 and Permanova+ v1.0.3 [32]. Briefly, square root transformed relative abundances were used to obtain a resemblance matrix based on the S17 Bray-Curtis similarity. The above matrix was then used into downstream analysis including hierarchical cluster analysis and non-metric multidimensional scaling (nMDS). Group centroids were determined from the above Bray-Curtis resemblance matrix and used to generate a new resemblance matrix of distances between groups. A network focusing on the high frequency OTUs detected in both vagina and uterus was generated with QIIME 1.8 and implemented in Cytoscape 3.2.0.

Pyrosequencing data was analysed with the Quantitative Insights Into Microbial Ecology (QIIME 1.8) [33]. Multiplexed sequences were assigned to samples based on their unique nucleotide barcode while any low quality or ambiguous reads were removed. In order to reduce the amount of erroneous Operational Taxonomic Units (OTUs), denoising of the dataset was performed using denoise_wrapper.py [34] in two hi1.4x large instances in EC2 Amazon Web Services. Chimeras were detected with ChimeraSlayer [35] and removed from the dataset. Datasets were deposited into the Sequence Read Archive (SRA) under accession numbers SRX3849466 and SRX3849984. *De novo* picking of OTUs, at 97% of sequence identity, was carried out with uclust [36]. Representative sequences were aligned to the best matching sequence in the Greengenes 13_8 core reference alignment using the PyNAST method [37]. Taxonomic affiliations were assigned with uclust and a phylogenetic tree constructed using FastTree [38]. Jackknifed-supported UPGMA trees of samples was constructed from rarefied OTU tables using UniFrac distances [39].

## Results

### Incidence of postpartum endometritis

In total, 26 out of 97 animals were diagnosed as healthy (26.8%) and 24 were diagnosed with postpartum endometritis (24.7%). The remaining 47 (48.5%) animals presented with metritis; short term, acute uterine disease characterised by the presence of purulent vaginal mucus on 7 and/or 14 DPP but that had resolved by 21 DPP. Given that the focus of the present work was on comparing the microbiomes of healthy cows and cows that developed postpartum endometritis, animals diagnosed with metritis were not further included. Samples that failed to produce PCR products or whose replicates produced highly variable TRFLP profiles were also excluded. Table 1 shows the number of samples remaining for healthy and endometritic cows at different times postpartum.

**Table 1.** Summary of vaginal and uterine sampling and clinical assignments*

| Days postpartum | Healthy |  | Endometritic |  |
| --- | --- | --- | --- | --- |
|  | Vagina | Uterus | Vagina | Uterus |
| Precalving** | 18 | - | 16 | - |
| 7 | 21 | 14 | 15 | 12 |
| 21 | 11 | 14 | 15 | 11 |
| 50 | 9 | 12 | 12 | 11 |

\* Vaginal and uterine swab samples were taken in duplicate on 7, 21 and 50 DPP.
\*\* Precalving samples were only taken from the vagina at -7 DPP.

### Comparison of vaginal and uterine microbiomes in the reproductive tract of healthy cows postpartum

With the aim of comparing vaginal and uterine microbiomes, duplicate vaginal and uterine swabs were collected from dairy cows on days 7, 21 and 50 postpartum (Table 1) and analysed by T-RFLP of the 16S rRNA gene. Overall, 311 OTUs were observed (Fig 1A) at different frequencies among the sampled animals (Fig 1B). Approximately 50% of the OTUs were observed in 5% or less of the cows, showing an important degree of variation among individuals. OTUs of medium representation, shared by 7 to 23% of the cows, constituted about 40% of the microbiome. The remaining 10% of the OTUs were detected in 23 to 68% of the animals. Vaginal and uterine samples produced similar T-RFLP profiles (Fig 1A), OTU distributions (Fig 1B) and shared highly represented OTUs, as visualised in an OTU network (Fig 1C). Unsurprisingly, ordination of samples by non-metric multidimensional scaling (nMDS) failed to form clusters by site of sampling (Fig 1D). In addition, the data failed to show a significant temporal variation in the microbial communities at different times postpartum (Permanova P = 0.087). Taken together, these results indicate that although there is a high degree of variation among individuals, a core microbiome exists in the postpartum reproductive tract, not only among different animals but also between the vaginal and uterine microbiomes as no significant differences were detected (Permanova P = 0.38).

**Figure 1.**
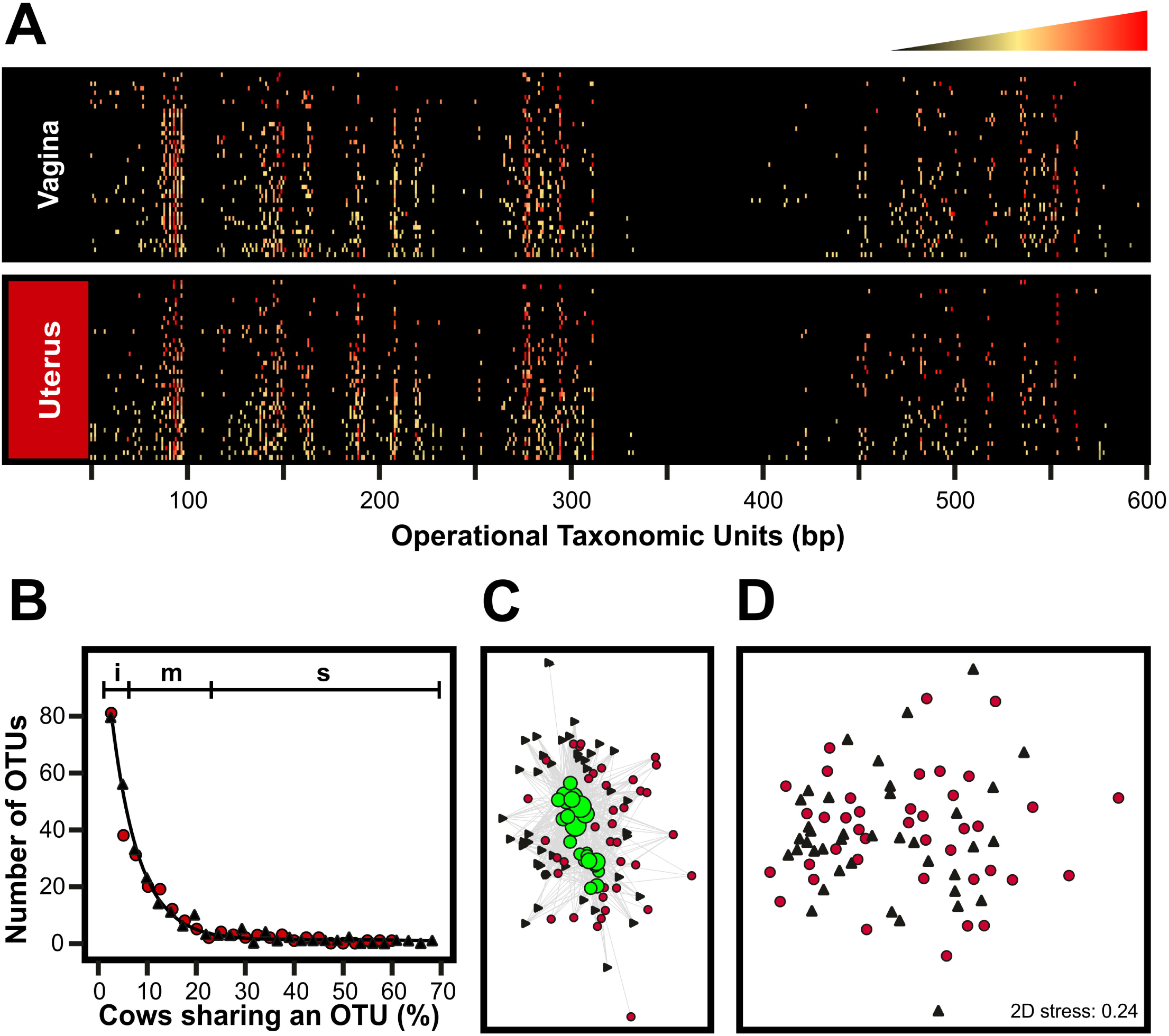
T-RFLP-based comparison of the vaginal and uterine microbiomes of healthy cows postpartum. PCR amplicons of the 16S rRNA were obtained from DNA samples using primers 6-FAM-labeled 27F and 1052R and digested with *Msp*I restriction enzyme. Fluorescently labeled terminal restriction fragments (TRFs) were resolved in an ABI 3730xl genetic analyzer (Applied Biosystems). GeneScan Liz 600 size standard was used for fragment sizing using GeneMapper v4.0 (Applied Biosystems). Operational Taxonomic Units (OTUs) were assigned in T-Align from TRFs representing at least 0.5% of the total signal and consistently found in duplicated samples. **A)** Heat map representing community profiles obtained from postpartum vaginal and uterine samples. The profile of each sample (rows) is populated by OTUs of varying lengths (columns). The relative abundance of the OTUs in a given profile is indicated by the colour key on the top where black and red represent the lowest and highest values and the midpoint (yellow) was set as the 95 percentile. The heat map was produced in Excel 2010 using the conditional formatting function and subsequently imported as image into CorelDraw X4. **B)** Distribution of OTUs in the reproductive tract of healthy cows postpartum. Black, vagina; Red, uterus. The regions marked on top show the contribution of OTUs to the total number of observed OTUS. i, 50%; m, 40%; s, 10%. **C)** Network showing high frequency OTUs (green circles) present in both vaginal (black triangles) and uterine (red circles) samples. OTUs and samples are connected through edges. The size of the green circles represents the OTU frequency among sampled animals. Samples with higher number of connections are displayed closer to the OTUs. **D)** Ordination of samples by non-metric multidimensional scaling (nMDS) based on the Bray-Curtis resemblance of the samples. Vaginal and uterine samples are represented by black triangles and red circles, respectively.

### Community changes associated with postpartum endometritis

This study hypothesises that the microbiome of the bovine reproductive tract is related to the reproductive health status of the animal and that changes in microbial community structure would be especially relevant in animals affected by postpartum endometritis. Comparison of T-RFLP datasets showed that the microbiome of the reproductive tract of cows that subsequently developed postpartum endometritis is significantly different to that of healthy cows (Permanova P = 0.001) and that the microbial populations change over time (Permanova P = 0.014). Furthermore, a change in community structure on 7 DPP is associated with the subsequent development of endometritis (Fig 2). The community change included both a decline in OTUs that were otherwise highly represented in healthy animals, as well as the appearance of a sub-community associated with postpartum endometritis that is observed in both vagina and uterus (Fig 2A and 2B). Hierarchical cluster analysis of group centroids separated the sample groups into two major clusters. Cluster 1 (Branch 1 in Fig 2C) contains both vaginal and uterine populations from cows developing postpartum endometritis at 7 DPP (V7E and U7E). Cluster 2 is formed by two branches (Branches 2a and 2b in Fig 2C). Branch 2a contains healthy cows at 7 DPP (V7C and U7C), uterine microbiomes of healthy cows at 21 DPP (U21C) and both vaginal and uterine microbiomes of endometritic cows at 21 and 50 DPP (V21E, U21E, V50E and U50E). Branch 2b contains the precalving groups (VPcC and VPcE), the group of vaginal microbiomes from healthy animals at 21 DPP (V21C), and both vaginal and uterine microbiomes at 50 DPP (V50C and U50C). Recovery of the community structure in cows that developed postpartum endometritis was evident at 21 DPP and continued at 50 DPP (Fig 2). Highly represented OTUs in healthy animals that were lost at 7 DPP in animals developing postpartum endometritis, started to reappear while endometritis-associated OTUs decreased (Fig 2A and Fig 2B). These results suggest a succession in the microbial communities as a consequence of the disturbance in the reproductive tract during calving. In addition, they show that the disturbance in healthy animals is lower, which may be associated with a faster clearance than in cows developing postpartum endometritis.

**Figure 2.**
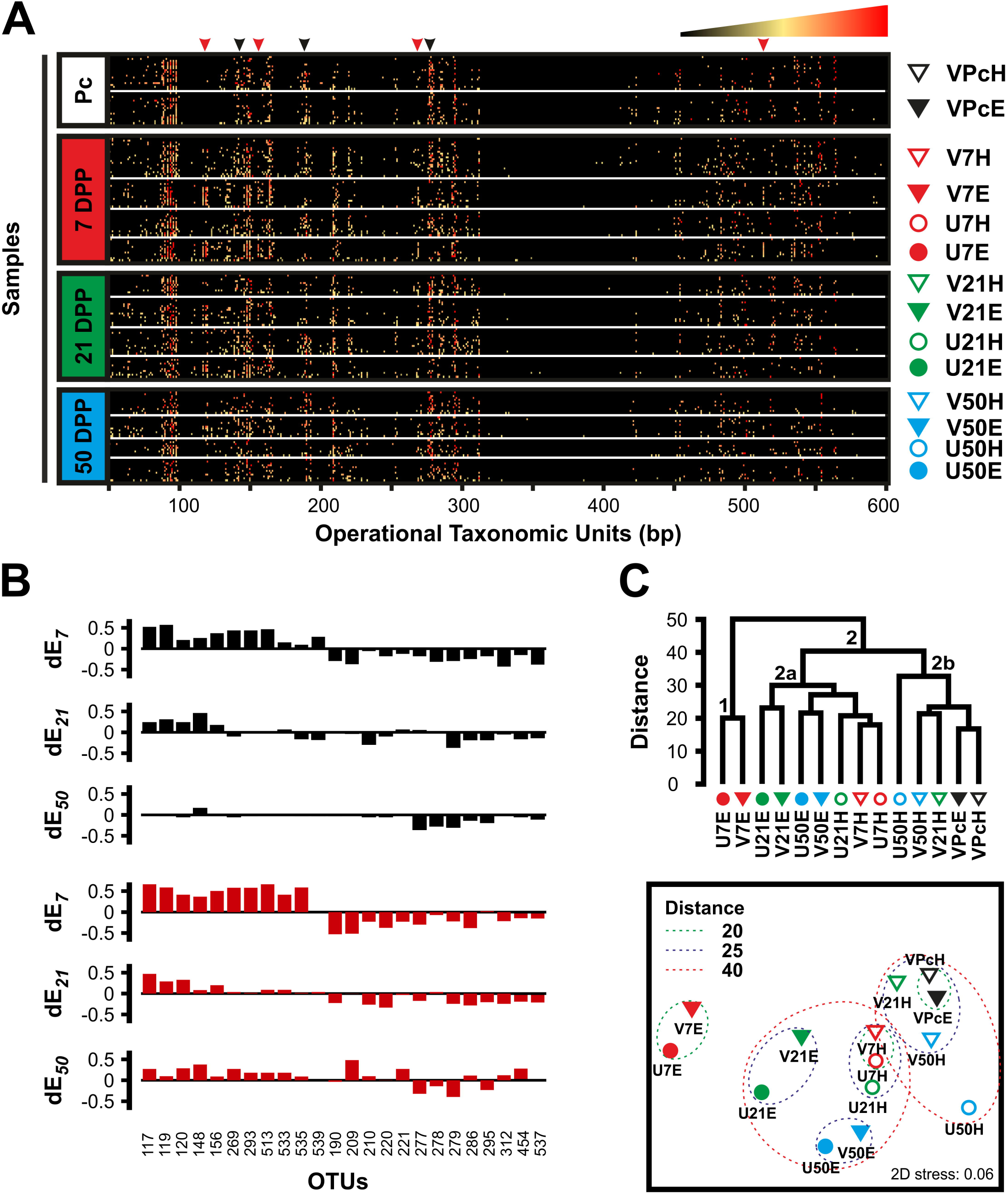
Temporal analysis of the microbiome associated with the reproductive tract of cows pre- and postpartum. **A)** T-RFLP-based analysis of the vaginal (triangles at the right of the heatmaps) and uterine (circles) microbiomes of healthy (empty symbols) and endometritic (solid symbols) cows pre-calving (Pc) and at different times postpartum (7, 21 and 50 DPP). The profile of each sample (rows) is populated by OTUs of varying lengths (columns). Arrowheads at the top of the heatmap indicate some of the changes associated with animals developing postpartum endometritis. Black arrowheads, loss of highly represented OTUs; Red arrowheads, appearance of OTUs. The relative abundance of the OTUs in a given profile is indicated by the colour key at the top right corner. Amplicons of the 16S rRNA were obtained from DNA samples by nested PCR and as described in Materials and Methods section. Fluorescently labeled terminal restriction fragments (TRFs) were resolved in an ABI 3730xl genetic analyzer. GeneScan Liz 600 size standard was used for fragment sizing using GeneMapper v4.0. OTUs were assigned in T-Align from TRFs representing at least 0.5% of the total signal and consistently found in duplicated samples. The heat maps were produced in Excel 2010 using the conditional formatting function. **B)** Differential OTUs in the reproductive tract of dairy cows postpartum. The dE*_t_* value accounts for the difference in frequency of a given OTU between endometritic and healthy cows (dE*_t_* = *f_e_* – *f_h_*) at a given time postpartum (denoted by *t*). Thus, the frequency of OTUs with positive values of dE is increased in endometritic animals while negative values show increased frequency in healthy cows at a given time postpartum. Left to right plots respectively correspond to 7, 21 and 50 DPP in vagina (Black plots) and uterus (Red plots). **C)** Complete linkage hierarchical cluster analysis (Top panel) and non-metric multidimensional scaling (Bottom panel) of groups. Distances are based on Bray-Curtis dissimilarity of group centroids. Each group is identified using the same symbols showed at the right of the heat maps in A. The analysis was performed in PRIMER6 and PERMANOVA+ with group centroids obtained from a resemblance matrix generated from square root transformed T-RFLP data. Figures were re-drawn using CorelDraw X4.

### Vaginal and uterine microbiomes are most similar in cows developing postpartum endometritis at 7 DPP

So far, we have shown that the most important changes in the community structure of the reproductive tract microbiome happen at 7 DPP (Fig 2), that there is a strong component of the microbiome associated with individual-specific OTUs (Fig 1B) and that vaginal and uterine share a core microbiome in healthy animals (Fig 1C and 1D). Regression analysis of shared OTUs in paired samples of vagina and uterus at 7 DPP provided evidence that the observed similarity is based not only on the presence of shared OTUs but also on their relative abundances (Fig 3A and 3B). The probability of detecting OTUs in a given microbiome due to neutral processes, such as bacterial dispersion, is proportional to the abundance of the same OTU in a source microbiome [40]. The results presented in Fig 3C and 3D show that the higher the relative abundance of shared OTUs in the vagina, the higher their frequency of detection in uterus. This is consistent with the occurrence of a neutral process that results from the homogenization of the vaginal and uterine microbiomes due to the disruption of the compartmentalization of the reproductive tract during parturition. In addition, higher coefficients of determination were observed in cows developing postpartum endometritis as compared with cows that achieved clearance suggesting a delayed differentiation of the vaginal and uterine microbiomes in the former group.

**Figure 3.**
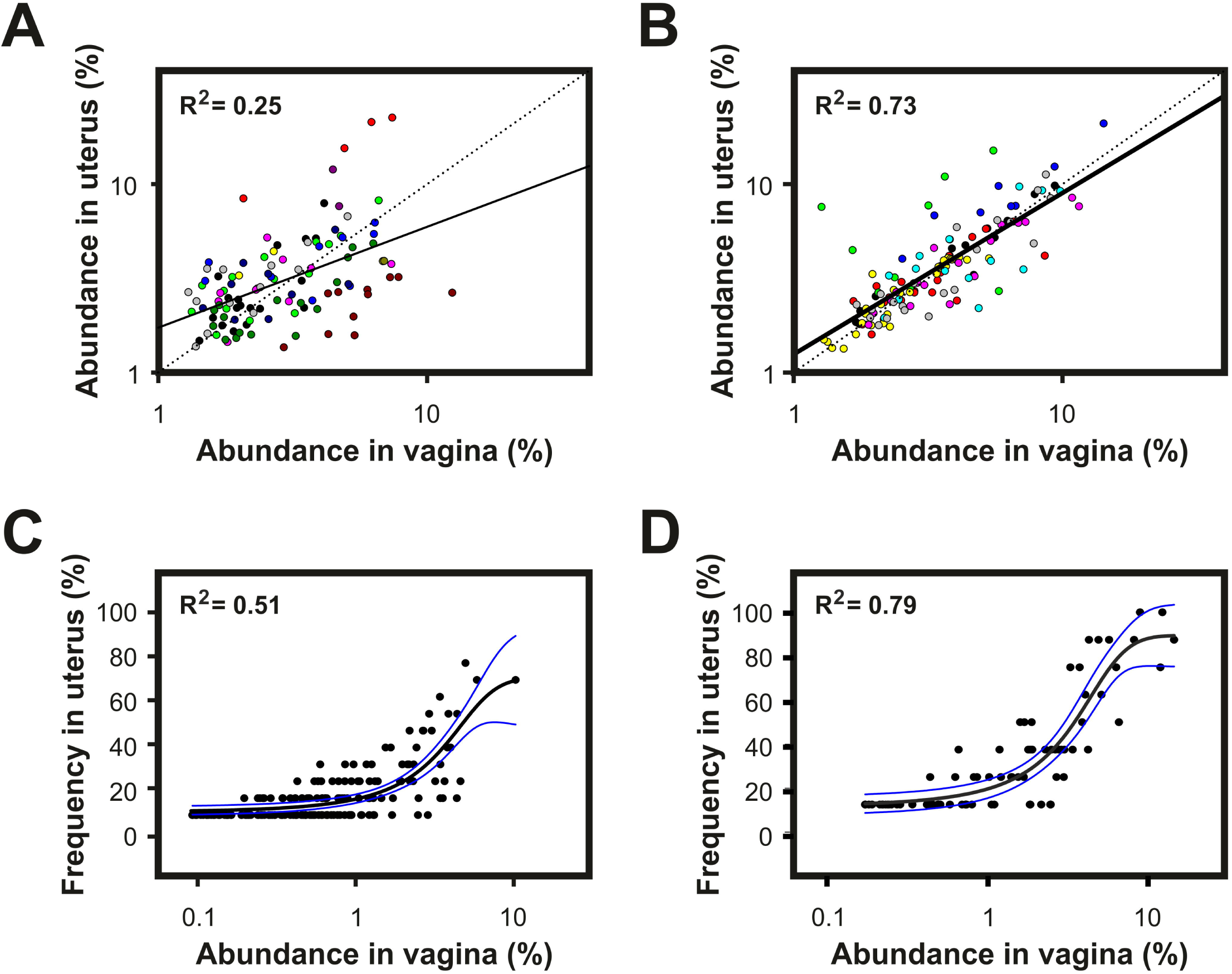
Disruption of the compartmentalization in the reproductive tract during parturition results in homogenization of vaginal and uterine microbiomes. **A and B)** Scatter plots of the abundance of OTUs observed in both vagina and uterus of dairy cows at 7 DPP. OTUs from each cow are displayed in different colour. The line of best fit (solid line) was obtained by least squares regression. The coefficient of determination shows the goodness of fit. The dotted line indicates the expected line assuming perfect correlation. **C and D)** Frequency of OTU detection in the uterus of dairy cows at 7 DPP as a function of their relative abundance in vagina. The relative abundance of a given OTU in the community of vagina was calculated as the average fluorescence signal associated to the OTU in the vagina of sampled animals. The observed frequency of detection for each OTU in uterus was calculated as number of cows in which the OTU was detected / total number of cows. Data was fitted to a 3-parameter sigmoid using the dynamic fit wizard of SigmaPlot 13. The higher the coefficient of determination the best the overall best-fit solution. **A and C)** Healthy cows; **B and D)** Cows developing postpartum endometritis.

In agreement with the above results, comparison of the similarity between paired vaginal and uterine microbiomes showed that, at 7 DPP, the vaginal and uterine microbiomes of cows developing postpartum endometritis are more similar than in animals that cleared the transient infection postpartum (61.9 ± 15.0% vs 25.7 ± 12.7%, P < 0.001) (Fig 4A). Differentiation between vaginal and uterine communities was evident from a decreased similarity in paired samples over time (38.9 ± 22.1 % at 21 DPP and 27.8 ± 13.1% at 50 DPP) (Fig 4B-4D). Taken together, these results suggest that both differences among individuals and the presence of shared OTUs in vagina and uterus mask differences in the composition of vaginal and uterine microbiomes that only become evident when comparing paired samples. In addition, they also suggest that the microbial succession in animals developing postpartum endometritis is delayed as compared with animals capable of clearing the infection postpartum, whose vaginal and uterine microbiomes differentiated as early as 7 DPP.

**Figure 4.**
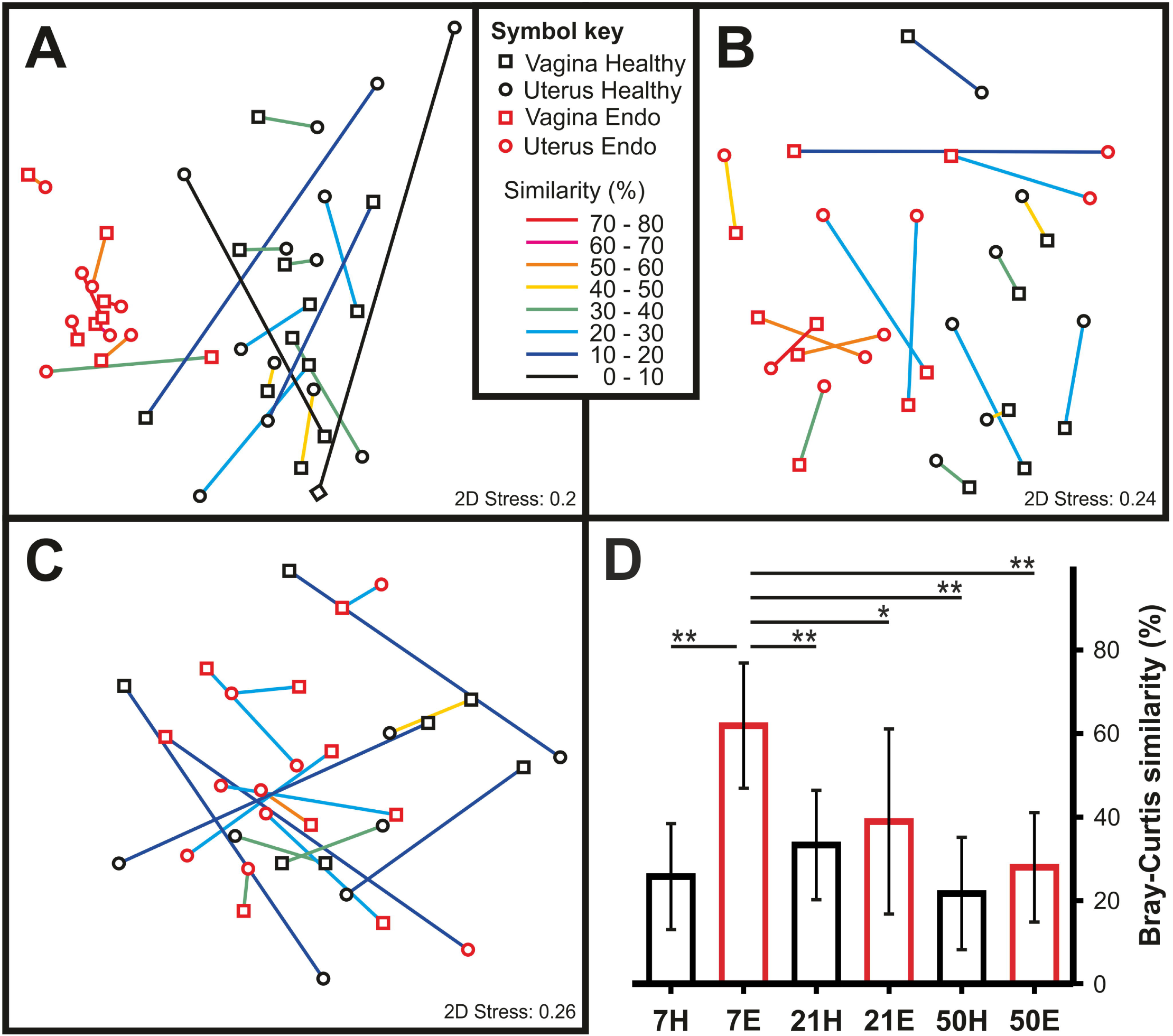
Differentiation of the vaginal and uterine microbiomes. **A-C)** Non-metric multidimensional scaling of vaginal (squares) and uterine (circles) samples collected at **A)** 7, **B)** 21 and **C)** 50 days postpartum from animals that cleared the transient infection (black symbols) or that developed postpartum endometritis (red symbols). Vaginal and uterine samples originating from the same animal are linked. The colour of the link represents the Bray-Curtis similarity between the microbiomes associated with each compartment of the reproductive tract. Links between paired samples were added in CorelDraw X4. **D)** Average and standard deviation of the Bray-Curtis similarities of paired vaginal and uterine samples at 7, 21 and 50 days postpartum from healthy (black bars) and endometritic cows (red bars). Asterisks above the horizontal lines show the presence of significant differences between groups: *, P < 0.05; **, P < 0.001.

The above results prompted the question whether the prepartum vaginal communities in dairy cows could differentiate between healthy animals and those that will develop endometritis. Comparison of T-RFLP datasets obtained from 35 cows indicated that the vaginal microbial community of the pre-calving cow is not related to postpartum uterine health (Permanova P = 0.7655) (Fig S1).

### Pyrosequencing of the vaginal microbiome of cows at 7 days postpartum

So far, the largest differences in the microbiomes of healthy cows and cows developing postpartum endometritis were observed at 7 DPP. Interestingly, those differences are contained in the vaginal microbiome. Thus, we decided to further analyse the vaginal microbiome of 30 cows, 20 healthy and 10 developing postpartum endometritis, by pyrosequencing an amplicon containing the variable regions v1 to v3 of the 16S rRNA. A dataset of 701189 high-quality, non-chimeric, sequences was obtained with an average of 23373 ± 6898 sequences per animal. A representative set was generated by clustering sequences in (OTUs) at 97% of identity. In total, 8504 non-chimeric OTUs were found with an average 933 ± 614 per sample. Table S2 shows a summary of the metrics for each sample. The Chao1 metric estimated that the average number of species was 1410 ± 860 per sample. The current sequencing effort was sufficient to cover 95.47 ± 3.56% of the species as determined by Good’s estimator of coverage. The diversity was observed in the range of *H’* 0.61 to 6.29 with an average of *H’* 3.82 ± 1.75 and the evenness ranging from *J’* 0.1 to 0.84 (Table S2). In agreement with the T-RFLP data, the wide range in the values of these metrics show an important difference in the communities associated with individual samples.

A total of 7576 out of 8504 OTUs had representative sequences in the Greengenes 13_8 database and were distributed into 21 phyla, 52 classes, 90 orders, 174 families and 379 genera. Overall, the six most abundant phyla were Firmicutes (64%), Bacteroidetes (27.7%), Fusobacteria (2.9%), Proteobacteria (1.8%), Tenericutes (1.3%) and Actinobacteria (1.1%). The remaining 928 OTUs were only assigned within the kingdom Bacteria. In spite of accounting for a large percentage of the OTUs, their contribution to the total abundance was relatively minor as only 4683 sequences (ie. 0.67% of the observations) were associated with these.

### Dysbiosis in the vaginal microbiome of cows developing postpartum endometritis

In agreement with the results yielded by T-RFLP, pyrosequencing data showed distinct microbiomes at 7 DPP between healthy cows and in cows subsequently developing postpartum endometritis. Analysis of principal coordinates showed that 33% of the variation of the data was explained by the first principal coordinate (PC1), which separated healthy from endometritic cows (Fig 5A). PC2, accounting for 17% of the variation, was most likely related to differences among individuals. These changed communities featured a significant reduction in the number of observed OTUs (P < 0.0001), bacterial diversity (*H’*, P < 0.0001) and species evenness (*J’*, P = 0.0003) as compared to healthy animals (Fig 5B and Table S2). The collapse of bacterial diversity was evident in a rarefaction curve where the number of observed species in cows that developed postpartum endometritis approached the asymptote much faster than cows that cleared the postpartum infection (Fig 5C). The total number of estimated species by the Chao1 metric was 649.6 ± 322.14 and 1789.95 ± 789.58 for endometritic and healthy cows, respectively (Fig 5A and Table S2).

**Figure 5.**
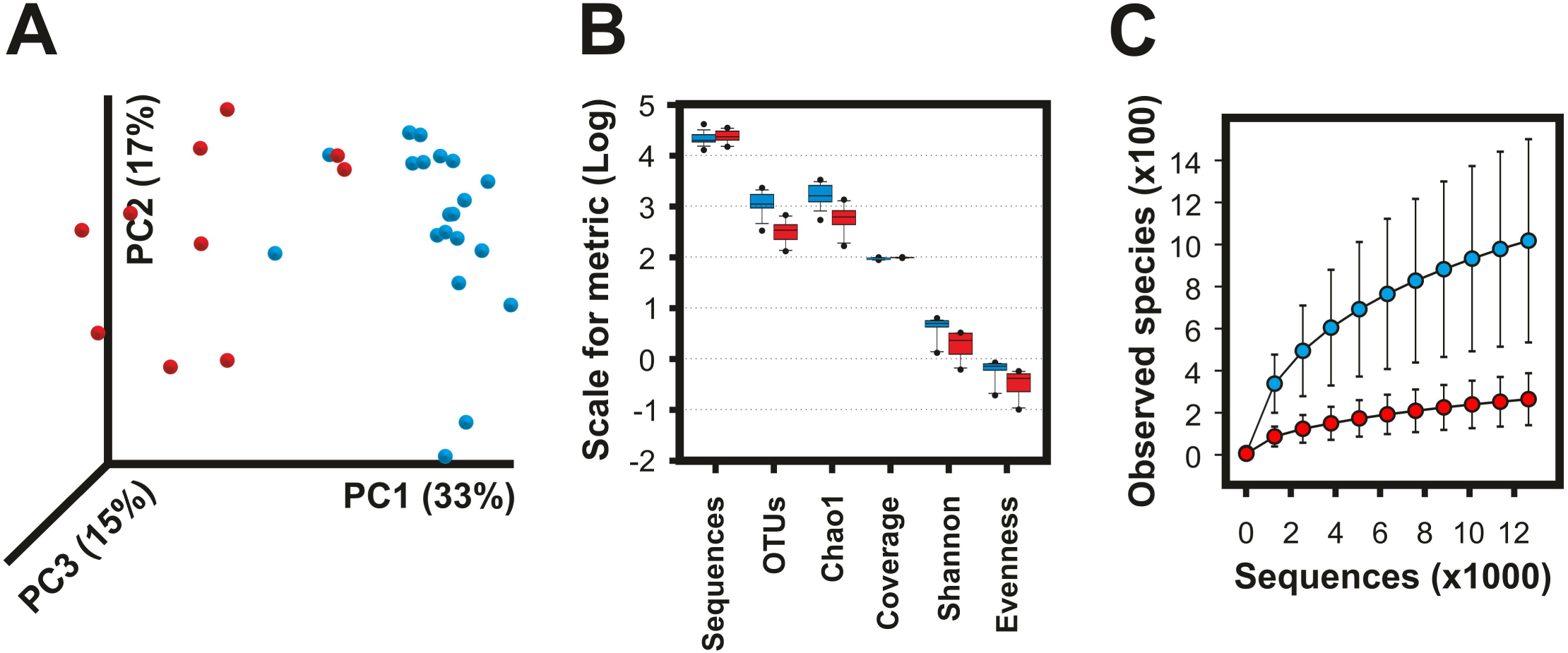
Collapse of the vaginal microbiome in cows developing postpartum endometritis. Differences in the vaginal microbiome of cows developing postpartum endometritis (red) and healthy cows (blue) were captured by 454 pyrosequencing of the v1-v3 16S rRNA at 7 days postpartum. OTUs at 97% of identity were generated in QIIME 1.8 using the pick_de_novo_otus.py pipeline. **A)** Principal components analysis showing a clear separation of samples by health status. **B)** Box plot summary of the diversity metrics of the vaginal microbiome showing lower richness and lower diversity in animals that developed postpartum endometritis (See Table S2). **C)** Rarefaction analysis of observed species. The curves represent the average of each group. Error bars are the standard error of the corresponding groups.

Taxonomic assignments of OTUs at phylum level showed that the microbiome of healthy animals displays high content of Firmicutes while most cows that developed postpartum endometritis had an increased representation of Bacteroidetes (Fig 6 and Fig S2A). The median Firmicutes to Bacteroidetes ratio (F/B) in the healthy group was 4.02 while cows that developed postpartum endometritis displayed a median F/B of 0.64.

### The vaginal microbiome displays different community types in both healthy and endometritic animals

Hierarchical cluster analysis using a weighted UniFrac metric resulted in four clusters of samples (Fig 6A). Cluster I is formed by 16 out of the 20 samples collected from healthy animals, whose microbiomes featured the larger number of observed OTUs in the dataset (1297.5 ± 584.4), high diversity (*H’* 5.25 ± 0.60) and evenness (*J’*, 0.75 ± 0.07) (Table 2). Cluster IV was formed by two healthy cows that displayed a microbiome with low diversity but intermediate species richness (*H’* = 1.66 and *J’* = 0.24, Table S2). The phylum Firmicutes constituted the 74.9 ± 7% and 95.1 ± 1 % of the total abundance of the microbiomes of respectively clusters I and IV (Fig 6A and Table 2). An important difference between clusters I and IV is that while the most abundant OTU in animals of cluster I constituted the 12.5 ± 7.5% of the microbiome, a single OTU accounted for more than 70% of the microbiomes of cows in cluster IV.

**Figure 6.**
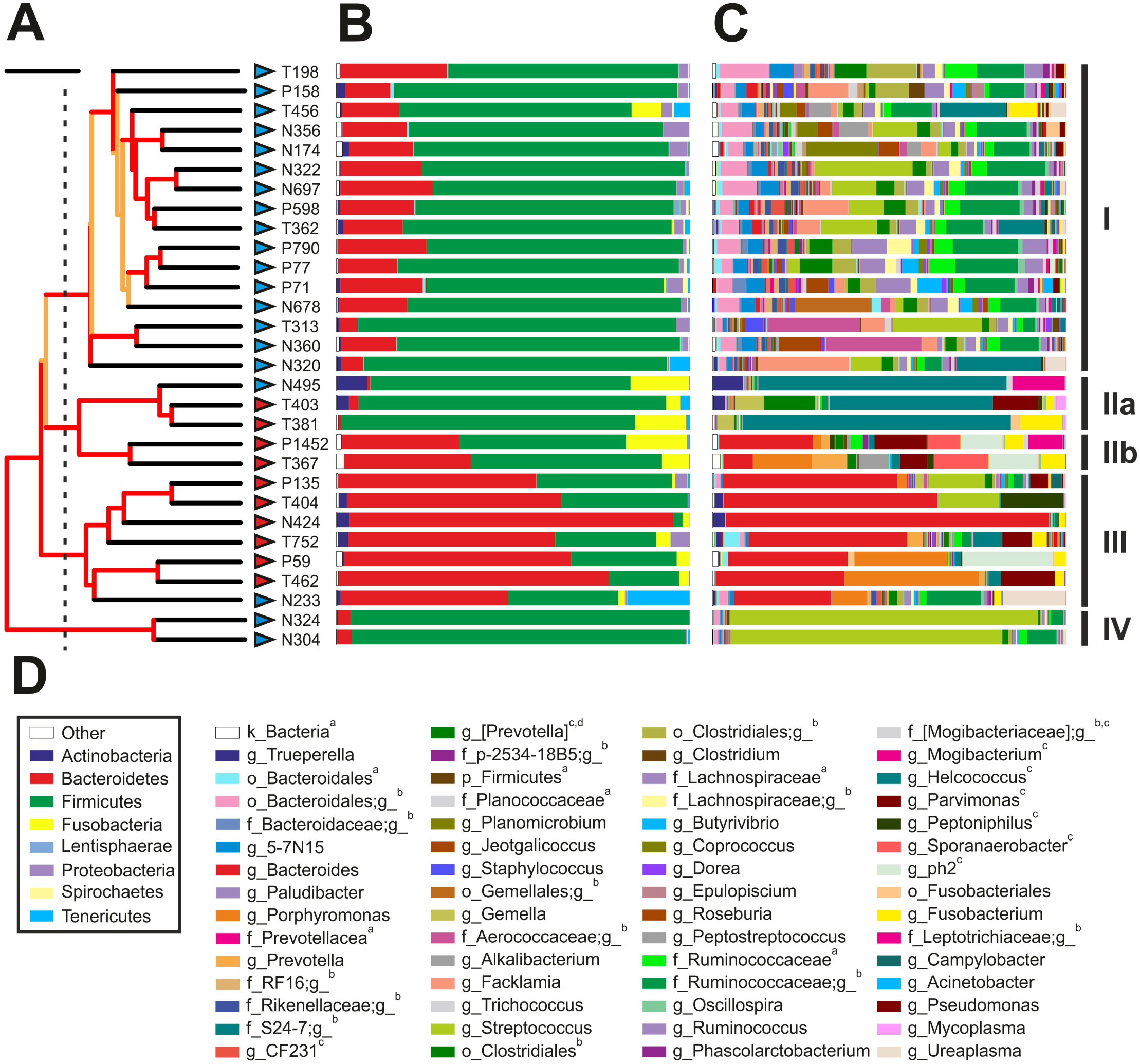
Taxonomic composition of the vaginal microbiome of cows at 7 DPP. **A)** Whole-microbiome weighted UniFrac hierarchical cluster. Jackknife support of internal tree nodes is colour coded. Red, 75-100%; orange, 50-75%. The identity of the samples is given to the right of the branch tips. Scale bar represents 0.1 distance. Cluster membership as defined by the branching pattern of this dendrogram is shown at the far right of the figure and is based upon a distance cut off displayed by a dashed line. **B) and C)** Taxonomic composition at phylum and genus levels, respectively. Each bar represents the complete microbiome as observed in each of the vaginal samples. **D)** Colour key of selected phyla (within box) and genera from B and C. Superscripts: **^a^**OTUs with ambiguous assignment below the indicated taxonomic level; **^b^**OTUs that although matching reference sequences in the Greengenes 13_8 database, no taxonomic name has been defined. In these cases, the lowest taxonomic name is provided; **^c^**OTUs matching reference sequences for which taxonomic changes above the rank of genus have been recommend by Greengenes based on whole genome phylogeny; **^d^**OTUs matching Genus name contested. Complete taxonomy plots derived from QIIME are shown as supporting information (Fig S3).

**Table 2.** Summary of metrics for clusters obtained by Weighted Unifrac Hierarchical analysis

| Cluster | Sequences | OTUs | $H'$ | $J'$ | Chao1 | Coverage | Characteristics |
| --- | --- | --- | --- | --- | --- | --- | --- |
| I | 20752.1<br>(4284.6)<sup>a</sup> | 1297.5**<br>(584.4) | 5.25***<br>(0.6) | 0.75***<br>(0.07) | 1901.6**<br>(843) | 93.4**<br>(3.5) | Healthy, high richness, high diversity |
| IIa | 27010.7<br>(15756.5) | 523<br>(160.7) | 1.63<br>(0.7) | 0.26<br>(0.11) | 776.6<br>(127.5) | 99.3<br>(0.3) | Endometritis, low richness, low diversity |
| IIb | 24998<br>(6137.7) | 319.5<br>(21.9) | 3.23<br>(0.04) | 0.56<br>(0.01) | 429.8<br>(35.6) | 99.7<br>(0.1) | Endometritis, low richness, high diversity |
| III | 26652.4<br>(6177.1) | 435.4<br>(325.3) | 2.28<br>(1.18) | 0.38<br>(0.16) | 755.9<br>(491) | 98.3<br>(1.5) | Endometritis, low richness, low diversity |
| IV | 25780.5<br>(9975.2) | 985.5*<br>(99.7) | 1.66<br>(0.51) | 0.24<br>(0.08) | 1413.6*<br>(270.8) | 95.9*<br>(1.2) | Healthy, high richness, low diversity |

<sup>a</sup> Standard deviations are given in brackets
\* $P < 0.05$ ; \*\* $P < 0.01$ ; \*\*\* $P < 0.0001$ indicate significantly different as compared to the same metric of non-starred clusters.

Cows that developed postpartum endometritis branched in clusters II and III. Interestingly, both clusters displayed similar metrics. In contrast to cluster I, both cluster II and III were characterised by a lower species richness and lower diversity (Table S2). Interestingly, the vaginal microbiomes of cows N495 and N233, both classified as healthy by inspection of the vaginal mucus score, were most similar to the microbiomes of cows that developed postpartum endometritis and branched in clusters II and III, respectively (Fig 6A and Table S2). The latter was characterised by high content of Bacteroidetes (65 ± 14.7%, F/B = 0.49) (Fig S2B). In contrast, cluster II was formed by two branches differing in the number of Firmicutes and Bacteroidetes. The Firmicutes to Bacteroidetes ratio branch IIb was low (F/B = 1.46) while samples in branch IIa had very low representation of Bacteroidetes and therefore displayed a F/B of 78.83. At the phylum level both cluster IIa and IIb had a high content of Fusobacteria (~12%) relative to other clusters (Fig S2B).

Firmicutes in cluster I comprised OTUs belonging to classes of Clostridia and Bacilli. A single order, Clostridiales, was represented in the first, whereas the second was represented by the orders of Lactobacillales (72.2%), Bacillales (21%) and Gemellales (6.2%). The most abundant taxon in cows of cluster I, formed by OTUs assigned to the family Ruminococcaceae, which accounted for 19.4 ± 7.4% of the abundance of the microbiome and 41.4% of Clostridia (Fig S2B). Moreover, in contrast to the low diversity in clusters IIa, III and IV, in which microbiomes were dominated by a single OTU, the combined abundance of the top five OTUs of the family Ruminococcaceae in cows of cluster I accounted for 17.1 ± 11.2% of the taxon. Although this taxon was found in 29 out of the 30 animals under study, its relative abundance was lower in endometritic cows of clusters IIa, IIb and III (P < 0.01). Interestingly, the representation of this taxon in cow N233, who was healthy but branched in cluster III was in similar abundance as in cows of cluster I, while OTUs affiliated to Ruminococcaceae in the two healthy cows of cluster IV constituted 4.5% and 11.5% of their microbiomes. Similar results were observed for the family of Lachnospiraceae. In addition, while single OTUs dominated the diversity of Bacteroidetes in cows that developed postpartum endometritis, healthy animals of cluster I displayed higher diversity and evenness indexes of Bacteroidetes than cows in clusters IIb and III (P < 0.001).

### The loss of diversity in vaginal microbiome of cows developing postpartum endometritis is characterised by the presence of dominant OTUs at genus level

At genus level, the two cows in cluster IV produced 26 and 34 OTUs of the genus Streptococcus, which comprised 76.8% and 87.2% of their microbiomes. From these, a single OTU in each cow dominated at least 98% of the representation of the above genus explaining the sharp decrease in bacterial diversity (Fig 6C and Fig S2B). The high content of Bacteroidetes in 5 out of the 7 cows of cluster III was due to the presence of a highly dominant OTU of the genus Bacteroides contributing 52.5 ± 24.2% of the total vaginal microbiome in cows that developed postpartum endometritis. In addition, OTUs of Porphyromonas constituted 26.3 and 38% of the microbiomes of two cows in cluster III (Fig 6C). Cluster IIa featured a high content of Firmicutes (>80%). However, in contrast to cluster I that showed high indexes of diversity and evenness, the microbiomes of cows in cluster IIa were dominated by OTUs of the family Tissierellaceae (69.9%), of which 96% were affiliated to the genus Helcococcus (Fig S2B). The phylum of Firmicutes in cluster IIb also displayed high proportion of Tissierellaceae (39.6%). However, in contrast to cluster IIa, genera ph2, Sporanaerobacter and Parvimonas were in similar proportions and constituted 90.8% of the above family. Another important characteristic of clusters IIa and IIb is the high content of Fusobacteria. The major OTUs within this phylum had close relatives from the family Leptotrichiaceae (99% identity) and genus Fusobacterium (100% identity). Similar to cluster III, the OTUs with largest abundance in cows of cluster IIb were Bacteroidetes from the genus Bacteroides and Porphyromonas, respectively.

## Discussion

This study revealed early signatures in the microbiome of cows that subsequently developed postpartum endometritis. T-RFLP analysis showed that these signatures were characterised by the appearance of a community associated with endometritis as well as the decline of OTUs highly represented in healthy animals. Significantly changed communities were evident at different time points between 7 DPP and 50 DPP. The occurrence of bacterial succession during the postpartum period was previously reported for bovine uterine microbiota [12,15,17]. Here, we show that this succession also includes the vaginal microbiota. The greatest differences in microbiome composition were observed at 7 DPP between cows that achieved uterine clearance and those that developed postpartum endometritis. Previous studies of either the vaginal [19–24] or uterine microbiome [11–18] have been reported but to date no study has compared the vaginal and uterine microbiomes in the same cows. Analysis of paired vaginal and uterine microbiomes at 7 DPP suggested the mixing of bacteria due to neutral processes during calving. Presented data also suggested a differentiation of the vaginal and uterine microbiomes that was most evident in cows developing postpartum endometritis.

Despite widespread use of vaginal mucus assessment for the clinical evaluation and classification of the reproductive health status of cows, the bovine vaginal microbiome has only recently become the focus of analysis [19–24]. Using pyrosequencing of an amplicon containing the V1 - V3 hypervariable regions of the 16S rRNA, we showed at 7DPP the presence of a complex microbiome in the vaginas of healthy cows and a dysbiotic microbiome in cows developing postpartum endometritis. High content of Firmicutes, high Firmicutes to Bacteroidetes ratio and a high diversity index were some of the most prominent features of the vaginal microbiome in healthy cows. A major reduction in the vaginal bacterial diversity of cows that subsequently developed postpartum endometritis was associated with an increased abundance of OTUs of *Bacteroides*, *Helcococcus*, and *Fusobacterium*, among other genera. Similar results were recently shown in the vagina of cows [21,23]. In agreement with our data, a recent study showed that at 7 DPP the number of Bacteroidetes is significantly higher in the vagina of cows that were subsequently diagnosed with endometritis at 35 days in milk (DIM) [24]. However, other studies have found different results. For example, in contrast to major changes in the microbiome of cows developing postpartum endometritis observed in this work, the most significant difference between metritic and healthy cows was an increased rate of isolation of *E. coli* in infected cows [22]. The authors suggested a lack of a stable microbiota in the bovine vagina and concluded that vaginal bacteria were likely contaminants from different sources, including skin, faeces and/or from the environment. Although our data supports the conjecture of an unstable (i.e. changing) microbiota during the postpartum period, the bias introduced by enrichment in culture-dependent approaches is well known, as only a very small fraction of the microbiome can be cultured on any given media and growth conditions [41]. A study using denaturing gradient gel electrophoresis (DGGE) and clone libraries of 16S indicated that there is a lower bacterial diversity in the vaginal microbiome of healthy cows as compared to cows with endometritis at 30-40 DPP. Dominant taxa included *Lactobacillus* and *Weissella* while endometritic animals did not show any dominant species [20]. Interestingly, while *Bacteroides*, *Prevotella*, and *Clostridium perfringens* strains were equally prevalent in healthy and endometritic cows, *Fusobacterium*, *Enterococcus* and *E. coli* were found in higher numbers in diseased animals as determined by qPCR [20]. Another study using Ion Torrent showed evidence of distinct communities in the healthy and diseased groups. *Bacteroides* and *Enterobacteriacea* were the largest taxa in both groups [19]. However, the number of sequence reads was highly dissimilar between groups: 31,000 and less than 1000 for the endometritic and healthy groups, respectively, making difficult any comparison between the microbiomes associated with groups of different health status. While some of the varying findings among the above studies may be related to the employment of different technologies, sampling times or other factors intrinsic to each study, our findings showed that postpartum endometritis may be associated with microbiomes of varying composition. In fact, our pyrosequencing analysis showed at least three different vaginal microbiome types associated with cows developing postpartum endometritis. Microbiome type IIa was dominated by OTUs of the genus *Helcococcus* while types IIb and III were characterised by a large content of *Bacteroides*. *Fusobacteria* was higher in vaginal microbiome types IIa and IIb than in type III and rare in healthy animals. Whether different microbiome types correspond to different types of postpartum endometritis or represent intermediate states of recovery into a healthy microbiome is yet to be determined.

In agreement with our T-RFLP data, suggesting an arrest of the differentiation of the vaginal and uterine microbiomes in cows developing postpartum endometritis, *Bacteroides* and *Fusobacterium* have been found to be most abundant in the uterine microbiome of cows with uterine infection at different times postpartum [11,14,16–18,42,43]. Sequencing of the V4 hypervariable region of the 16S rRNA showed the progression of the uterine microbiome of dairy cows during the first 6 DPP [17]. A rapid succession resulted in a shift from Proteobacteria to Bacteroidetes and Fusobacteria as the most abundant phyla in the uterus of metritic cows. Similar to our findings in vagina, diseased cows displayed lower uterine bacterial richness and diversity indices related to an increased abundance of OTUs of *Bacteroides, Porphyromonas* and *Fusobacterium* [17,18]. Failure to cure metritis, either spontaneously or with antibiotic treatment, was related to increased relative abundances of the above genera and a corresponding decrease in bacterial diversity in uterus [18]. In a study relying on DGGE and analysis of clone libraries, the uterine fluid of two healthy and two metritic cows at 10 DPP showed that the most abundant OTUs in metritic cows belonged to the phylum Fusobacteria followed by Bacteroidetes [11]. Another study based on pair-ended MiSeq sequencing of an amplicon containing the V1 and V2 hypervariable regions of the 16S DNA of uterine fluids obtained from cows with pyometra, slaughtered at no less than 22 DPP, found that the five most abundant OTUs in uterine fluids belonged to families of Fusobacteriaceae, Bacteroidaceae, Pasteurellaceae and Porphyromonadaceae [16]. A higher prevalence of *Bacteroides* and *Fusobacterium* was also reported upon pyrosequencing analysis of V1 and V2 hypervariable regions of the microbiota of uterine lavages of cows with severe endometritis at 35 DIM as compared to either the healthy group or to cows with mild endometritis [14]. In another study, Ruminococcaceae, Bacteroidaceae, and an unclassified family that belonged to class Bacteroidia were the three most abundant families in endometrial biopsies from healthy cows at 4 weeks postpartum (WPP) as well as from both healthy and endometritic cows at 7 WPP [15]. Interestingly, our pyrosequencing analysis showed that Ruminococcaceae and Bacteroidaceae were the top one and six most abundant families in the vaginal microbiome of healthy cows at 7 DPP. In addition, the T-RFLP data showed that the differentiation between vaginal and uterine communities of endometritic cows was evident from a decreased similarity in paired samples. Taken together, our data and those of others are consistent with a bacterial succession in the reproductive tract in which the differentiation of vaginal and uterine microbiomes towards the recovery of their native states are conducive to reproductive health and the achievement of a new pregnancy. Conversely, a delayed differentiation of vaginal and uterine microbiomes is in line with impaired uterine clearance, decreased conception rates and lower success of first service pregnancy rates in the endometritic cow [44]. Interestingly, high prevalence of *Bacteroides*, *Ureaplasma*, *Fusobacterium* and *Arcanobacterium* were still observed in the uterine microbiome of cows that failed to become pregnant after 200 DIM [14], strengthening the link between a severe arrest in the differentiation of the uterine microbiome and poor fertility.

Bacterial infection and tissue damage is a normal event occurring in the postpartum period. Involution of the postpartum uterus is a highly regulated process in which an early inflammatory response early postpartum is followed by a stage of proliferation and repair [2,3]. Comparison of RNAseq profiles of the endometrium at 7 and 21 DPP unveiled that the above transition was arrested in cows with cytological endometritis [27]. Sustained inflammation was also observed in the endometrium of cows with postpartum endometritis in the same time frame [45]. The observed collapse in the diversity of the vaginal microbiome of endometritic cows at 7 DPP and the arrested differentiation of vaginal and uterine microbiomes in cows developing postpartum endometritis are in line with those data. A likely scenario implies different metabolic landscapes in the reproductive tract of healthy and endometritic cows early after calving, resulting from their distinct microbiotas. Support for the above scenario comes from recent metagenomic analyses of the microbiome of the bovine uterus where significant differences in the repertoire of functional gene categories were observed in healthy and metritic cows within 3 and 12 DIM [42] and cows with purulent vaginal discharge between 25 and 35 DPP [43]. These studies showed that adhesins, bacteriocins and antibacterial peptides and tolerance to colicin E2 are produced only by the uterine microbiota of healthy cows early postpartum and continues until at least 35 DPP. In contrast, the uterine microbiota of metritic and PDV cows appears to change from cold shock and acid stress in the former to increased modification of lipid A and production of toxins in the later [42]. Changes in the composition of the microbiome, and therefore in the metabolic landscape of the reproductive tract, may impact uterine involution as a result of the dysregulated inflammatory response induced by the presence of highly abundant bacteria activating specific signalling pathways and thus causing a different pattern of endometrial gene expression [46]. This in turn is likely to affect the transition from the inflammatory to proliferation and repair stage of the postpartum uterus observed in healthy cows [2,27]. It is difficult to establish causality between dysbiosis and inflammation [47]. Thus, it is not clear if changes in the composition of the microbiome precede the inability of the cow to regulate the immune response or whether both the microbiome and the dysregulated inflammatory response are factors predisposing the onset of postpartum endometritis. In any case, a synergistic effect may occur where failure of each of these factors exacerbates the other. In other words, failure to clear a highly changed microbiome characterised by a low bacterial diversity and dominated by few bacterial taxa may set an innate immune response in overdrive. Alternatively, an excessive inflammatory response may contribute to the differential elimination of bacterial species, allowing the overgrowth of bacteria able to avoid the innate immune response.

## Acknowledgements

This study was funded by Science foundation Ireland (07/SRC/B1156) and the Department of Agriculture, Food and the Marine (13–S–472). The funding agencies played no role in the study design, data collection and analysis, decision to publish, or preparation of the manuscript. The authors acknowledge the contribution of farmers who participated in this study. We thank Fiona Carter, Maria Benson and Carlotta Sacchi for providing technical assistance.

## Supporting information

**Table S1. Dataset of Terminal Restriction Fragment Length Polymorphism.** (DOCX)

**Table S2. Diversity metrics of the vaginal microbiome of dairy cows at 7 days postpartum.** (DOCX)

**Figure S1. Hierarchical cluster analysis of pre-calving dairy cows based of their vaginal microbiomes.** The microbiomes associated with vaginal samples obtained from cows before calving were compared in a resemblance matrix based on the Bray-Curtis similarity. The health status of each cow was assessed depending on the outcome of the transient postpartum infection. The tips of the branches are colour coded according to the outcome of postpartum health status: Black, non-susceptible; Red, susceptible to postpartum endometritis. Community profiles were determined by T-RFLP of the 16S rRNA as described in the section of Materials and Methods. Analysis was performed in PRIMER6 and the figure was re-drawn in CorelDraw X4. (TIF)

**Figure S2. Category-based taxonomic composition of the vaginal microbiome of cows at 7 DPP.** Taxonomic composition at phylum and genus levels, respectively. Each bar represents the average of the vaginal microbiome in each of the following categories: **A)** Clinical assignment. h, healthy; e, endometritis **B)** Cluster as defined in Figure 6A, **C)** Farm of collection. The colour key of selected phyla (within box) and genera from A, B and C is placed at the right of the figure. Superscripts: **^a^**OTUs with ambiguous assignment below the indicated taxonomic level; **^b^**OTUs that although matching reference sequences in the Greengenes 13_8 database, no taxonomic name has been defined. In these cases, the lowest taxonomic name is provided; **^c^**OTUs matching reference sequences for which taxonomic changes above the rank of genus have been recommend by Greengenes based on whole genome phylogeny; **^d^**OTUs matching Genus name contested. (TIF)

**Fig S3. Summary of the taxonomic composition of the vaginal microbiome of cows at 7 DPP.** (HTML) Original output generated by QIIME. To visualise it double click on bar_charts.html.

## References

1. Sheldon IM, Williams EJ, Miller AN, Nash DM, Herath S (2008) Uterine diseases in cattle after parturition. Vet J 176: 115–121.

2. Foley C, Chapwanya A, Creevey CJ, Narciandi F, Morris D, Kenny EM, Cormican P, Callanan JJ, O’Farrelly C, Meade KG (2012) Global endometrial transcriptomic profiling: transient immune activation precedes tissue proliferation and repair in healthy beef cows. BMC Genomics 13: 489.

3. Chapwanya A, Meade KG, Foley C, Narciandi F, Evans AC, Doherty ML, Callanan JJ, O’Farrelly C (2012) The postpartum endometrial inflammatory response: a normal physiological event with potential implications for bovine fertility. Reprod Fertil Dev 24: 1028–1039.

4. Sheldon IM, Cronin J, Goetze L, Donofrio G, Schuberth HJ (2009) Defining postpartum uterine disease and the mechanisms of infection and immunity in the female reproductive tract in cattle. Biol Reprod 81: 1025–1032.

5. Williams EJ, Fischer DP, Pfeiffer DU, England GC, Noakes DE, Dobson H, Sheldon IM (2005) Clinical evaluation of postpartum vaginal mucus reflects uterine bacterial infection and the immune response in cattle. Theriogenology 63: 102–117.

6. Olson JD, Ball L, Mortimer RG, Farin PW, Adney WS, Huffman EM (1984) Aspects of bacteriology and endocrinology of cows with pyometra and retained fetal membranes. Am J Vet Res 45: 2251–2255.

7. Elliott L, McMahon KJ, Gier HT, Marion GB (1968) Uterus of the cow after parturition: bacterial content. Am J Vet Res 29: 77–81.

8. Griffin JF, Hartigan PJ, Nunn WR (1974) Non-specific uterine infection and bovine fertility. I. Infection patterns and endometritis during the first seven weeks post-partum. Theriogenology 1: 91–106.

9. Williams EJ, Fischer DP, Noakes DE, England GCW, Rycroft A, Dobson H, Sheldon IM (2007) The relationship between uterine pathogen growth density and ovarian function in the postpartum dairy cow. Theriogenology 68: 549–559.

10. Leblanc SJ, Osawa T, Dubuc J (2011) Reproductive tract defense and disease in postpartum dairy cows. Theriogenology 76: 1610–1618.

11. Santos TM, Gilbert RO, Bicalho RC (2011) Metagenomic analysis of the uterine bacterial microbiota in healthy and metritic postpartum dairy cows. J Dairy Sci 94: 291–302.

12. Santos TM, Bicalho RC (2012) Diversity and succession of bacterial communities in the uterine fluid of postpartum metritic, endometritic and healthy dairy cows. PLoS ONE 7: e53048.

13. Peng Y, Wang Y, Hang S, Zhu W (2013) Microbial diversity in uterus of healthy and metritic postpartum Holstein dairy cows. Folia Microbiologica 58: 593–600.

14. Machado VS, Oikonomou G, Bicalho ML, Knauer WA, Gilbert R, Bicalho RC (2012) Investigation of postpartum dairy cows’ uterine microbial diversity using metagenomic pyrosequencing of the 16S rRNA gene. Vet Microbiol 159: 460–469.

15. Knudsen LRV, Karstrup CC, Pedersen HG, Angen O, Agerholm JS, Rasmussen EL, Jensen TK, Klitgaard K (2016) An investigation of the microbiota in uterine flush samples and endometrial biopsies from dairy cows during the first 7 weeks postpartum. Theriogenology 86: 642–650.

16. Knudsen LR, Karstrup CC, Pedersen HG, Agerholm JS, Jensen TK, Klitgaard K (2015) Revisiting bovine pyometra--new insights into the disease using a culture-independent deep sequencing approach. Vet Microbiol 175: 319–324.

17. Jeon SJ, Vieira-Neto A, Gobikrushanth M, Daetz R, Mingoti RD, Parize ACB, de Freitas SL, da Costa ANL, Bicalho RC, Lima S, Jeong KC, Galvão KN (2015) Uterine microbiota progression from calving until establishment of metritis in dairy cows. Applied and Environmental Microbiology.

18. Jeon SJ, Lima FS, Vieira-Neto A, Machado VS, Lima SF, Bicalho RC, Santos JE, Galv+úo KN (2018) Shift of uterine microbiota associated with antibiotic treatment and cure of metritis in dairy cows. Vet Microbiol 214: 132–139.

19. Rodrigues NF, Kastle J, Coutinho TJ, Amorim AT, Campos GB, Santos VM, Marques LM, Timenetsky J, de Farias ST (2015) Qualitative analysis of the vaginal microbiota of healthy cattle and cattle with genital-tract disease. Genet Mol Res 14: 6518–6528.

20. Wang J, Sun C, Liu C, Yang Y, Lu W (2016) Comparison of vaginal microbial community structure in healthy and endometritis dairy cows by PCR-DGGE and real-time PCR. Anaerobe 38: 1–6.

21. Laguardia-Nascimento M, Branco KMGR, Gasparini MR, Giannattasio-Ferraz S, Leite LR, Araujo FMG, Salim ACdM, Nicoli JR, de Oliveira GCa, Barbosa-Stancioli EF (2015) Vaginal Microbiome Characterization of Nellore Cattle Using Metagenomic Analysis. PLoS One 10: e0143294.

22. Wang Y, Ametaj BN, Ambrose DJ, Ganzle MG (2013) Characterisation of the bacterial microbiota of the vagina of dairy cows and isolation of pediocin-producing Pediococcus acidilactici. BMC Microbiol 13: 19.

23. Wang Y, Wang J, Li H, Fu K, Pang B, Yang Y, Liu Y, Tian W, Cao R (2018) Characterization of the cervical bacterial community in dairy cows with metritis and during different physiological phases. Theriogenology 108: 306–313.

24. Bicalho MLS, Santin T, Rodrigues MX, Marques CE, Lima SF, Bicalho RC (2017) Dynamics of the microbiota found in the vaginas of dairy cows during the transition period: Associations with uterine diseases and reproductive outcome. Journal of Dairy Science 100: 3043–3058.

25. Sheldon IM, Lewis GS, LeBlanc S, Gilbert RO (2006) Defining postpartum uterine disease in cattle. Theriogenology 65: 1516–1530.

26. Sheldon IM, Noakes DE, Rycroft AN, Pfeiffer DU, Dobson H (2002) Influence of uterine bacterial contamination after parturition on ovarian dominant follicle selection and follicle growth and function in cattle. Reproduction 123: 837–845.

27. Foley C, Chapwanya A, Callanan J, Whiston R, Miranda-CasoLuengo R, Lu J, Meijer W, Lynn D, O’ Farrelly C, Meade K (2015) Integrated analysis of the local and systemic changes preceding the development of post-partum cytological endometritis. BMC Genomics 16: 811.

28. Frank JA, Reich CI, Sharma S, Weisbaum JS, Wilson BA, Olsen GJ (2008) Critical evaluation of two primers commonly used for amplification of bacterial 16S rRNA genes. Appl Environ Microbiol 74: 2461–2470.

29. Heuer H, Krsek M, Baker P, Smalla K, Wellington EM (1997) Analysis of actinomycete communities by specific amplification of genes encoding 16S rRNA and gel-electrophoretic separation in denaturing gradients. Appl Environ Microbiol 63: 3233–3241.

30. Wang Y, Qian PY (2009) Conservative Fragments in Bacterial 16S rRNA Genes and Primer Design for 16S Ribosomal DNA Amplicons in Metagenomic Studies. PLoS One 4: e7401.

31. Smith CJ, Danilowicz BS, Clear AK, Costello FJ, Wilson B, Meijer WG (2005) T-Align, a web-based tool for comparison of multiple terminal restriction fragment length polymorphism profiles. FEMS Microbiol Ecol 54: 375–380.

32. Clarke K, Gorley R (2006) PRIMER v6: User Manual/Tutorial. PRIMER-E, Plymouth.

33. Caporaso JG, Kuczynski J, Stombaugh J, Bittinger K, Bushman FD, Costello EK, Fierer N, Pena AG, Goodrich JK, Gordon JI, Huttley GA, Kelley ST, Knights D, Koenig JE, Ley RE, Lozupone CA, McDonald D, Muegge BD, Pirrung M, Reeder J, Sevinsky JR, Turnbaugh PJ, Walters WA, Widmann J, Yatsunenko T, Zaneveld J, Knight R (2010) QIIME allows analysis of high-throughput community sequencing data. Nat Meth 7: 335–336.

34. Reeder J, Knight R (2010) Rapidly denoising pyrosequencing amplicon reads by exploiting rank-abundance distributions. Nat Meth 7: 668–669.

35. Haas BJ, Gevers D, Earl AM, Feldgarden M, Ward DV, Giannoukos G, Ciulla D, Tabbaa D, Highlander SK, Sodergren E, Methe B, DeSantis TZ, Petrosino JF, Knight R, Birren BW (2011) Chimeric 16S rRNA sequence formation and detection in Sanger and 454-pyrosequenced PCR amplicons. Genome Res 21: 494–504.

36. Edgar RC (2010) Search and clustering orders of magnitude faster than BLAST. Bioinformatics 26: 2460–2461.

37. Caporaso JG, Bittinger K, Bushman FD, DeSantis TZ, Andersen GL, Knight R (2010) PyNAST: a flexible tool for aligning sequences to a template alignment. Bioinformatics 26: 266–267.

38. Price MN, Dehal PS, Arkin AP (2010) FastTree 2 − Approximately Maximum-Likelihood Trees for Large Alignments. PLoS One 5: e9490.

39. Lozupone C, Knight R (2005) UniFrac: a New Phylogenetic Method for Comparing Microbial Communities. Applied and Environmental Microbiology 71: 8228–8235.

40. Venkataraman A, Bassis CM, Beck JM, Young VB, Curtis JL, Huffnagle GB, Schmidt TM (2015) Application of a Neutral Community Model To Assess Structuring of the Human Lung Microbiome. mBio 6: e02284–14.

41. Torsvik V, GoksØyr J, Daae FL (1990) High diversity in DNA of soil bacteria. Appl Environ Microbiol 56: 782–787.

42. Bicalho MLS, Machado VS, Higgins CH, Lima FS, Bicalho RC (2017) Genetic and functional analysis of the bovine uterine microbiota. Part I: Metritis versus healthy cows. Journal of Dairy Science 100: 3850–3862.

43. Bicalho MLS, Lima S, Higgins CH, Machado VS, Lima FS, Bicalho RC (2017) Genetic and functional analysis of the bovine uterine microbiota. Part II: Purulent vaginal discharge versus healthy cows. Journal of Dairy Science 100: 3863–3874.

44. Crowe MA, Williams EJ (2012) TRIENNIAL LACTATION SYMPOSIUM: Effects of stress on postpartum reproduction in dairy cows. J Anim Sci 90: 1722–1727.

45. Adnane M, Chapwanya A, Kaidi R, Meade KG, O’Farrelly C (2017) Profiling inflammatory biomarkers in cervico-vaginal mucus (CVM) postpartum: Potential early indicators of bovine clinical endometritis? Theriogenology 103: 117–122.

46. Carneiro LC, Cronin JG, Sheldon IM (2016) Mechanisms linking bacterial infections of the bovine endometrium to disease and infertility. Reprod Biol 16: 1–7.

47. Zhu W, Winter MG, Byndloss MX, Spiga L, Duerkop BA, Hughes ER, Büttner L, de Lima Romão E, Behrendt CL, Lopez CA, Sifuentes-Dominguez L, Huff-Hardy K, Wilson RP, Gillis CC, Tükel Ç, Koh AY, Burstein E, Hooper LV, Bäumler AJ, Winter SE (2018) Precision editing of the gut microbiota ameliorates colitis. Nature 553: 208.

